# Daily physical activity behavior: a compensatory factor to physical adaptations variability following a 12-week power training in older men

**DOI:** 10.64898/2026.09.18.752400

**Authors:** Layale Youssef, Haroun El-Oueslati, Justine Persouyré, Charlotte Pion, Paula Lago, Marc Bélanger, Mylène Aubertin-Leheudre

## Abstract

Physical activity (PA) is effective to counteract age-related declines. However, inter-individual variability in training adaptations may limit prediction of individual responses and intervention optimization. Daily PA behavior may contribute to this variability. The objective was to examine daily PA behavior through a 12-week supervised power training (PT) and its associations with clinical adaptations in 36 older men. Time of PA, step count, total (TEE) and active (AEE) energy expenditure, metabolic equivalents of task (METs), and sedentary time were assessed using a 3D accelerometer for three days surrounding PT sessions (Day-PT-1, Day-PT and Day-PT+1) at pre-(T0) and mid-intervention (6-week; T6). Physical performance, body composition and muscle function were assessed pre-(T0) and post-(T12) intervention. PA behavior outcomes didn’t change between T0 and T6, except for METs which decreased by 0.2 METS (T0:1.44±0.34 vs. T6:1.25±0.18; *p*=0.003 [13%]). At T6, day-to-day fluctuations were observed for PA behavior on Day-PT-1 and Day-PT+1 compared with Day-PT (all *ps*<0.05). Following the intervention, body composition and muscle characteristics improved significantly. Between T6-Day-PT-1 and T6-Day-PT, greater increases in time of PA were associated with lower Fast Timed Up and Go speed adaptation (*p*=0.02), and greater reductions in step count were associated with greater reductions in muscle power (*p*=0.02). Finally, increases in step count and time of PA were associated with greater upper limb strength (*p*=0.04) and muscle pennation angle (*p*=0.02). PA behavior appears modulated during a 12-week PT intervention in healthy older men. Thus, PA behavior surrounding training sessions could be an important compensatory factor contributing to adaptations variability.

## 1. Introduction

Aging is accompanied by a progressive decline in muscle mass, strength, and neuromuscular function (Clark, 2019) which contribute to the development of sarcopenia (Goodpaster et al., 2006). These alterations are leading to functional incapacities, higher risk of falls and fractures, and loss of independence in older adults (Dos Santos et al., 2017). In addition, it is also known that age-related changes in physical inactivity exacerbate these conditions (Shur et al., 2021), highlighting the importance of specific interventions to preserve musculoskeletal health and physical performance with advancing age.

Exercise is widely recognized as one of the most effective strategies to counteract these age-related declines (Izquierdo et al., 2021). Among various exercise modalities, power training, which combines force production with high contraction velocity, is recognized as a particularly effective approach for improving daily functional activities in older populations (Hazell et al., 2007). Indeed, power training specifically targets muscle power, a key determinant of mobility and daily function that declines more rapidly than maximal strength with aging (Porter & Metabolism, 2006). Previous studies have demonstrated that power training can induce significant improvements in functional parameters in older adults (Dulac et al., 2021; Izquierdo & Cadore, 2014; Jiménez-Lupión et al., 2023).

Despite power training’s benefits, substantial inter-individual variability in physiological adaptations is consistently observed (Chrzanowski-Smith et al., 2020), suggesting that factors beyond the prescribed exercise stimulus may influence the magnitude of training responses. One potential contributor to this variability could be daily physical activity behavior, which encompasses all movement performed outside of structured exercise sessions. While exercise interventions typically focus on prescribed training sessions, individuals spend the rest of their day engaged in habitual activities of varying intensity. Advances in wearable technology, particularly accelerometry, have enabled the objective and continuous assessment of these behaviors, providing detailed information on step count, energy expenditure, metabolic equivalent of task (METs) and sedentary time (Gorman et al., 2014; Lewis et al., 2017). These accelerometer-derived measures offer valuable insight into real-world activity patterns and represent an important, yet often overlooked, component of total daily physical activity (Godhe et al., 2022; Keadle et al., 2017). Importantly, physical activity behavior is often summarized as averaged values over several days like in weekly averages (Migueles et al., 2017), which may obscure meaningful day-to-day variability. Evidence from accelerometry studies indicates substantial within-individual variability in daily physical activity patterns (Bussmann & van den Berg - Emons, 2013;

Matthews et al., 2012). Such averaging can lead to the conclusion that behavior remains unchanged over time and may overlook fluctuations occurring across specific days, such as those before or after exercise sessions. Indeed, studies examining physical activity behavior in response to structured exercise have yielded inconsistent findings, with some reporting changes in activity or compensation outside exercise sessions, while others have found no significant changes in overall non-exercise physical activity (Castro et al., 2017; Fedewa et al., 2017; Mansfeldt & Magkos, 2023). This inconsistency may partly reflect differences in how physical activity is assessed and summarized, as aggregated measures may obscure day-to-day variations in activity and sedentary behavior. Examining behavior at a finer temporal resolution may therefore provide a more comprehensive understanding of behavioral responses to exercise and their potential contribution to the heterogeneity of adaptations observed following exercise interventions.

Therefore, the objective of the present study was to examine daily physical activity behavior during a 12-week power training intervention and its associations with clinical parameters in healthy community-dwelling older men. Specifically, the focus was placed on physical activity behavior on the days surrounding training sessions and explored its possible associations with physiological and clinical adaptations following the intervention. We hypothesized that: 1) daily physical activity would decrease on the days surrounding training sessions, with a greater reduction expected on the day following training, reflecting behavioral compensation associated with recovery from exercise-induced fatigue and muscle soreness, and a lesser but still meaningful reduction on the day preceding training, potentially reflecting behavioral adjustments to conserve energy and optimize performance during the upcoming session and; 2) some aspects of daily physical activity behavior (i.e., METs/d) would contribute to the variability in training-induced responses, given that habitual physical activity may influence the capacity to elicit further adaptations, particularly in healthy populations.

## 2. Methods

### Study design

A*-posteriori* per-protocol analysis was conducted using data from a randomized controlled trial (ClinicalTrials.gov Identifier: NCT03393650). The study protocol was approved by the UQAM Ethics Committee (CIEREH – UQAM; No.: A-120006), and all participants provided informed consent prior to enrollment in the assessments and intervention. The study design is presented in Figure 1.

**Figure 1.**
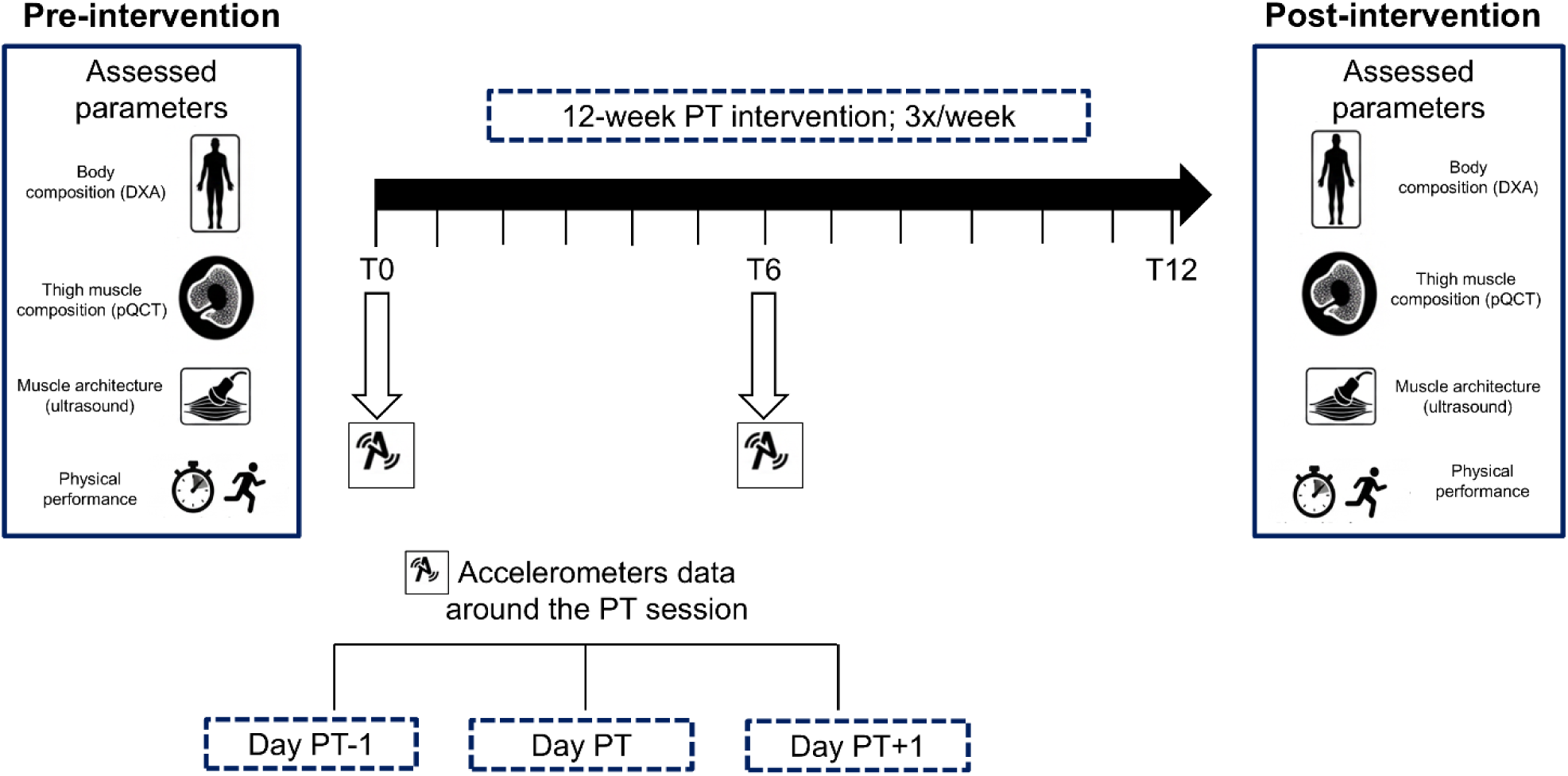
Study design. Legend: Study design over the 12-week intervention. Body composition parameters were assessed before and after the intervention. Symbols in the rectangles indicate the assessment methods used at each time point: DXA and pQCT (upper row) and ultrasound and physical performance parameters (lower row). PT = power training; DXA = dual-energy X-ray absorptiometry; pQCT = peripheral quantitative computed tomography. Pre = before the intervention; Post = after the intervention; T0 = baseline (pre-intervention); T6 = mid-intervention (week 6); Day PT = day of the power training session; Day PT-1 = the day before the power training session; Day PT+1 = the day after the power training session.

### Population

To be included in the main study, men participants need to be aged over 75 years old, in healthy status (non-smoker/ non-drinker; no major chronic condition; no unstable medication prescription; no metabolic disorder) but non physically active (less than 120 min/ week of exercise; not engaged in structured exercise training), cognitively able to consent, and able to follow a 12-week power training intervention (PARQ: Physical Activity Readiness Questionnaire). Recruitment was conducted within the community through flyers and workshops organized in collaboration with senior community centers in the Greater Montreal area.

For this a-posteriori analysis, participants were included if they had completed the pre- and post-intervention assessment and had valid accelerometer recordings at pre- (T0) and mid- (at 6 weeks) intervention for the following outcomes: total PA time (min/d), total energy expenditure (TEE; kcal/d), active energy expenditure (AEE; kcal/day), number of step (n/d), metabolic equivalent of task (METs/day), and sedentary time (min). To be considered as valid, records from the validated 3-axial accelerometer [(armband sensewear©; (Lopez et al., 2018)] need to have consecutive data from at least one day preceding the training session (Day PT-1), one day of training session (Day PT), and one day following the training session (Day PT+1). Based on these a-posteriori inclusion criteria, the final sample consisted of 36 participants with their characteristics presented in Table 1.

**Table 1.** Characteristics of the participants

|  |  |
| --- | --- |
| Sample size (n) | 36 |
| Sex (M; %) | 100 |
| Age (years) | 78.4 ± 10.9 |
| MoCA | 27.2 ± 2.1 |
| BMI (kg.m<sup>-2</sup>) | 26.3 ± 3.2 |
| Number of steps (n/day) | 8761 ± 4123 |
| Normal walking speed (m/s) | 1.47 ± 0.24 |

Legend: Data are presented as mean ± SD. n = number of participants; M= men; BMI = body mass index; MoCA = Montreal Cognitive Assessment

### Outcomes

#### -Anthropometric and body composition

Body mass (kg) was measured using a body composition monitor (Tanita BC-558©, Japan), and standing height (cm) was assessed using a stadiometer (Seca©, USA). Body mass index (BMI) was calculated as body mass divided by height squared (kg/m²).

Body composition parameters, including total, android and gynoid fat mass (FM) as well as total and appendicular (lower limb + upper limb) lean mass (LM), were assessed using dual-energy X-ray absorptiometry (DXA; GE Prodigy Lunar,© Madison, WI, USA). Fat mass was expressed as a percentage of total body mass (%) and lean mass were expressed as absolute values (kg).

#### - Muscle composition

Muscle composition of the right thigh was assessed using peripheral quantitative computed tomography [pQCT; Stratec XCT3000 system (STRATEC Medizintechnik GmbH©; Division of Orthometrix] at 33% of the femoral length. Scanning parameters, including voxel size (0.5 mm) and scanning speed (10 mm/s), were entered into the acquisition software. Image quality was subsequently evaluated visually by a second evaluator using a previously established movement artifact grading scale, with all scans receiving a quality score ≤3 (Blew et al., 2014). Image processing was conducted using the peripheral quantitative computed tomography density distribution plugin for BoneJ (version 1.3.11) (Doube et al., 2010), an open-source analysis tool. The precision error for muscle area, muscle density, subcutaneous adipose tissue area, and intramuscular adipose tissue area has been reported to range from 2.1-3.7%, 0.7-1.9%, 2.4-6.4%, and 3-42%, respectively (Frank-Wilson et al., 2015).

#### - Muscle architecture

Muscle architecture was evaluated using portable ultrasound system (Sonosite© Inc.). To ensure consistency between assessments, the measurement was assessed at the distal third of the femur and in a seated position with the knee flexed at 90° as previously suggested (Narici et al., 2004). Three ultrasound images of the vastus lateralis (VL) muscle were acquired using a 38-mm, 7.5-MHz linear probe. The probe was placed perpendicular to the skin surface and aligned with the muscle’s longitudinal axis. Muscle architecture parameters were obtained in the sagittal plane through the ImageJ software and included pennation angle, muscle thickness, and fascicle length (Narici et al., 2004).

#### - Muscle Function

*<u>Upper limb muscle strength</u>:* was assessed using a hand dynamometer (Lafayette©, USA). Participants were instructed to stand upright with the shoulder in a neutral position and the elbow fully extended while exerting maximal grip force for at least 4 s. Three alternating trials were performed with each hand, and the highest value (N) obtained was used for analysis. Relative Upper limb strength was calculated by normalizing absolute upper limb strength to body mass (Upper limb strength/ BW: N/kg).

*<u>Lower-limb muscle strength</u>:* was evaluated using the one-repetition maximum (1-RM) test performed on a seated leg press machine. Participants completed progressive loading attempts, with up to five repetitions per attempt, until the maximal load that could be lifted once was identified. A 4-min recovery period was provided between attempts. Absolute lower-limb muscle strength was expressed as the maximal load achieved (LLMS: kg).

*<u>Lower-limb muscle power</u> :*was estimated through the 10s sit-to-stand test. Using the validated takaï equation (Takai et al., 2009).

#### - Physical Performance

Participants’ physical performance was evaluated using components of the validated Senior Fitness Test (Rikli & Jones, 2013), including walking speed (self-paced 4 meter walking test), gait parameter (usual and fast 3-meter Timed Up-and-Go (TUG)), cardiorespiratory endurance (alternate-step), lower limb function (10 rep Sit to Stand) as well as mobility and endurance (6 minutes walking test). These tests have been chosen as they are related to mobility, physical fitness, risk of fall, morbidity and mortality in older adults. More specifically:

*<u>Usual walking speed:</u>* to assess usual walking speed (m/sec), participants began from a standing position at a designated starting line and walked 6 m at their usual pace following a verbal signal. They were instructed to complete the distance without assistance. The time to perform the first and the last meter were not recorded in the walking speed to avoid the acceleration and deceleration phases.

*<u>Gait parameter:</u>* the validated usual 3-m TUG was used to assess gait parameter. To perform the test, participants need to stand up from a chair without using their arms, walk at usual-pace 3-meter, turn around, return to the chair, and sit down. The total time required to complete the sequence was recorded. The procedure was subsequently repeated with participants instructed to walk the distance as quickly as possible safely and without running.

*<u>Lower limb function:</u>* the validated 10 repetitions sit to stand test (STS) was used to assess lower limb function. To perform the test, participants need to perform 10 sit to stand from a chair without using their arms.

*<u>Cardiorespiratory endurance:</u>* the validated 20-sec alternate-step test was used to assess cardiorespiratory function. To perform the test, participants need to alternately place the entire right and left foot onto the 20-cm-high step and return to the starting position. Participants were instructed to perform as many complete alternating movements as fast as possible within 20 s.

*<u>Mobility:</u>* the validated six-minute walking test was used to assess mobility. Participants were instructed to walk at their own pace along a flat, enclosed 30-m track to cover as much distance as possible in six minutes. They were permitted to stop and rest as needed. The distance covered was recorded at each minute and at the end of the test.

#### Training protocol

The power training (PT) intervention consisted of a 12-week program performed in groups of eight, three times on non-consecutive days per week in a laboratory facility, under the supervision of two kinesiologists (i.e. to ensure the safety, load and speed of each exercise).

Briefly, each training session included a 10-min warm-up and cool-down (stretching) and a main core of training divided in 2 parts. The first part included a set of four high-velocity resistance exercises targeting major muscle groups (leg curl, chest press, lateral pulldown, and seated leg press) and performed at 80% of the one-repetition maximum (1-RM) with a 1-0-2-0 tempo (i.e. to perform the concentric phase of each movement as rapidly as possible while maintaining a controlled eccentric phase). A 1-min recovery interval was provided between sets. To adjust loading, the 1-RM assessment was performed at weeks 2 and 6 and the resistance was also increased by 5% when participants could successfully complete three sets of 12 repetitions. The second part consisted of three sets of 10 repetitions of six functional exercises (wall squats using a Swiss ball, pelvic rotations on a Swiss ball, shoulder external rotations, stair climbing, dumbbell biceps curls, and the bird-dog exercise). Exercise difficulty was progressively modified according to instability level and participants perceived fatigue using the 10- point Borg scale.

### Statistical analysis

Normality of variables included in the correlation analyses was assessed using the Shapiro-Wilk test. Using the *afex* R package, repeated-measures analyses of variance (ANOVA) were performed to examine the change in physical activity behavioral parameters. The models included TIME [Pre- (T0) vs. Mid- (T6)] and DAY (Day PT-1 vs. Day PT vs. Day PT+1) as within-subject factors, as well as their interaction (TIME × DAY). When significant effects were observed, post-hoc pairwise comparisons were performed using estimated marginal means from the *emmeans* R package, with Bonferroni adjustment applied for multiple comparisons. T-tests were applied to evaluate changes in clinical parameters between pre- (T0) and post- (T12) intervention using the *tidyr* R package. Absolute changes (Δ) were calculated for physical activity behavior parameters as Mid - Pre and for clinical parameters as Post - Pre. Pearson correlations were used to assess the associations between changes in physical activity behavior parameters and changes in clinical parameters. Correlation analyses were performed using the *Hmisc* R package. The normality of the variables included in the correlation analyses was assessed using the Shapiro-Wilk test. All statistical analyses were conducted in R software (version 4.2; R Foundation for Statistical Computing, Vienna, Austria) and statistical significance was set at *p* < 0.05.

## 3. Results

### Physical activity behavioral parameter through the intervention and around the training session

Total physical activity time (Table 2 and Figure 2A): At T6, total physical activity time was lower on both the day preceding (PT-1) and the day following (PT+1) training compared with the training day (PT), withPre reductions of 54 min (38%) and 49 min (35%), respectively (*p*=0.0007, *d*=−0.744 and *p*=0.001, *d*=0.700). In contrast, no differences between days were observed at T0. When the three days were averaged, total physical activity time did not differ between T0 and T6 (*p*=0.513, *d*=0.111; Table S1), indicating that the mid-intervention changes were specific to the days surrounding training rather than reflecting an overall change in physical activity.

**Figure 2.**
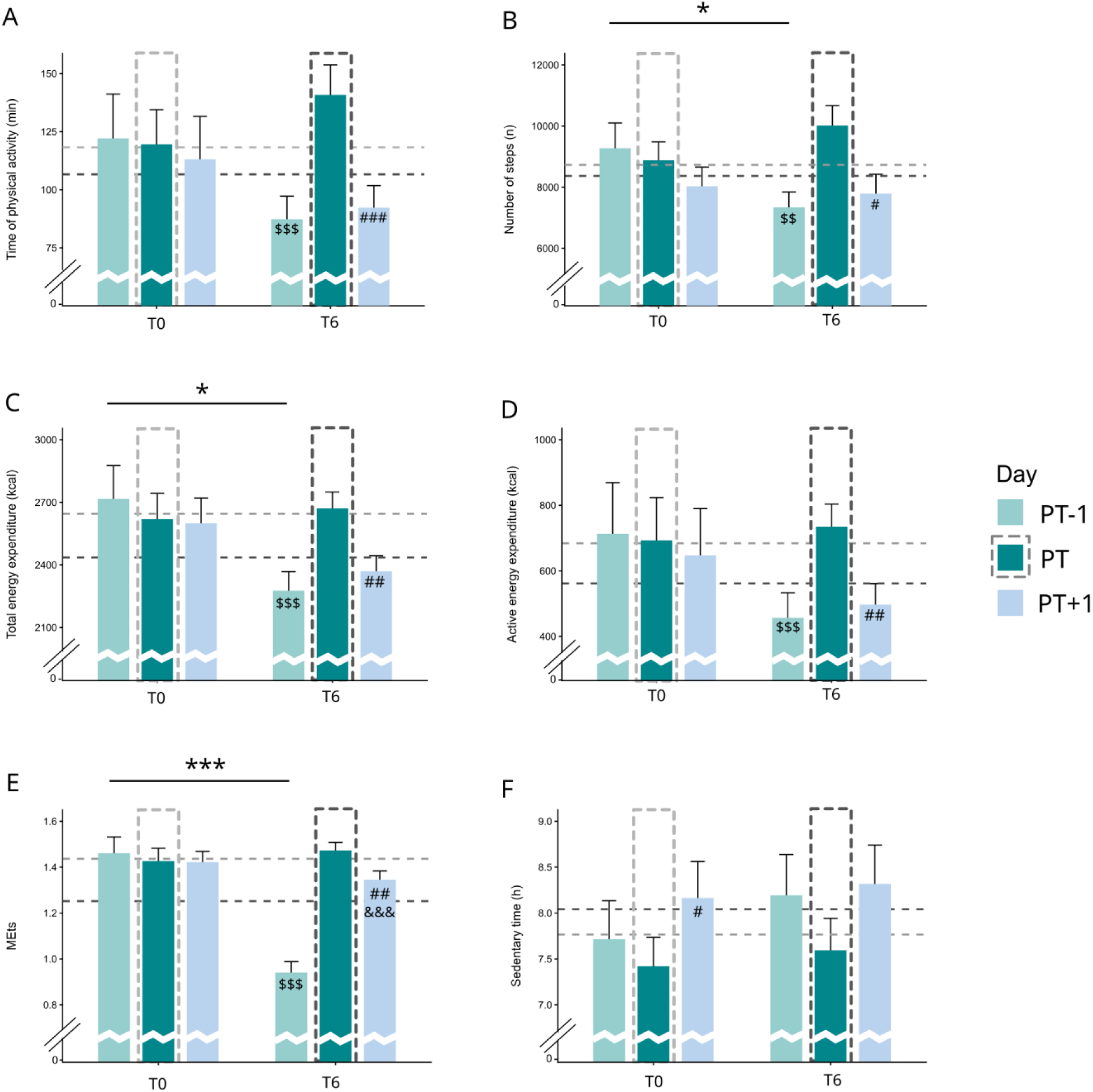
Changes in physical behavior parameters across the training days between Pre- and Mid-intervention. Legend: (A) Time of physical activity; (B) number of steps; (C) total energy expenditure; (D) active energy expenditure; (E) METs; (F) sedentary time. Dark green refers to the power training day (PT), while light green and light blue refer to the day before (PT-1) and after (PT+1) the training day, respectively. T0 = pre-intervention; T6 = mid-intervention corresponding to 6 weeks. Physical behavior parameters were assessed using a validated 3D accelerometer (armband sensewear©). min = minutes; kcal = kilocalories; METs = metabolic equivalents; h = hours. Error bars represent the standard error of the mean (SEM). Horizontal dashed lines represent the mean value across the three days (PT-1, PT, and PT+1) at each time point, with light grey representing T0 and dark grey representing T6. Dashed bars highlight the Day PT at each time point, with light grey representing T0 and dark grey representing T6. Statistical analyses were performed using two-way repeated measures ANOVA followed by post hoc comparisons. Symbols indicate significant pairwise differences within the same timepoint: $ = Day PT vs. Day PT-1; # = Day PT+1 vs. Day PT; & = Day PT-1 vs. Day PT+1.; * indicates a significant difference for a specific day between T0 and T6. For each symbol, one, two, or three repetitions indicate *p* < 0.05, *p* < 0.01, and *p* < 0.001, respectively.

**Table 2.**
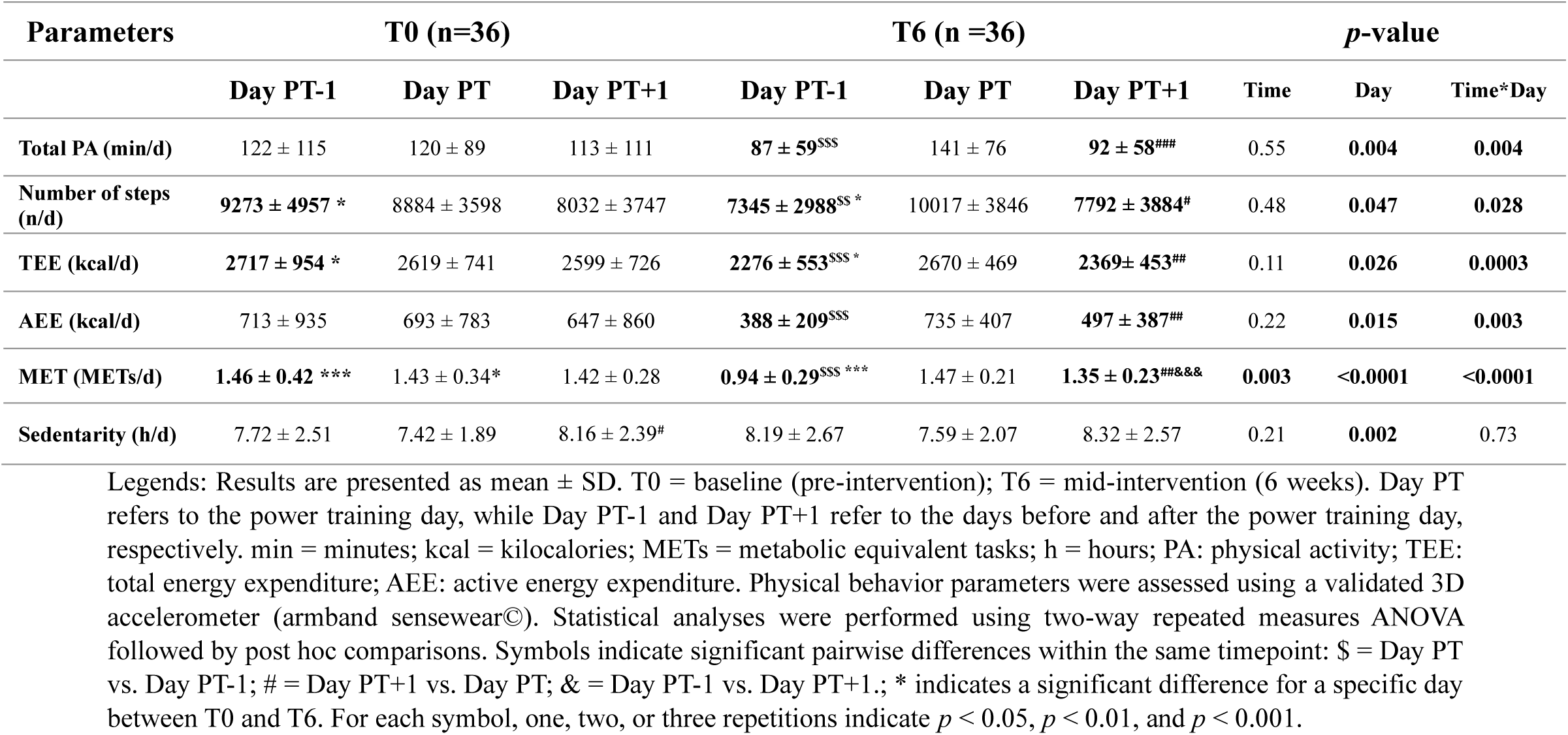
Changes in physical behavior parameters across the training days between pre- (T0) and mid-intervention (T6)

Number of steps (Table 2 and Figure 2B): Changes in the number of steps followed a pattern like that observed for PA time. At T6, participants took significantly fewer steps on Day PT-1 [-2672 steps (-27%)] and Day PT+1 [-2225 steps (-22%)] compared with Day PT (*p*=0.006, *d*=−0.586 and *p*=0.025, *d*=0.488, respectively). No differences between these days were observed at T0. In addition, the number of steps on Day PT-1 was significantly lower at T6 than at T0 (*p*=0.037, *d*=0.379). Despite these day-specific changes, the average number of steps across the three days did not differ between T0 and T6 (*p*=0.451, *d*=0.127).

Total energy expenditure (Table 2 and Figure 2C): Changes in TEE followed a pattern like that observed for the number of steps, although the magnitude of change was smaller. At T6, TEE was lower on Day PT-1 [-394 kcal (-15%)] and Day PT+1 (-301 kcal (-11%)) compared with Day PT (*p*=0.0003, *d*=−0.766 and *p*=0.001, *d*=0.673, respectively), whereas no differences between days were observed at T0. TEE on Day PT-1 was also significantly lower at T6 than at T0 (*p*=0.017, *d*=0.427). However, averaging TEE across the three days revealed no significant difference between T0 and T6 (*p* =0.103, *d*=0.279; Table S1).

Active energy expenditure (Table 2 and Figure 2D): The pattern of changes in AEE was like that observed for PA time. At T6, AEE was lower on Day PT-1 [-347 kcal (-47%)] and Day PT+1 [-238 kcal (-32%)] compared with Day PT (*p*=0.0001, *d*=−0.884 and *p*=0.004, *d*=0.644, respectively), whereas no differences between days were observed at T0. However, averaging AEE across the three days revealed no significant difference between T0 and T6 (*p*=0.434, *d*=0.134; Table S1).

Metabolic equivalent of task (Table 2 and Figure 2E): Overall, changes in METs followed a pattern similar to that observed for TEE. At T6, METs were lower on Day PT-1 (-0.53 (-36%)) and Day PT+1 (-0.12 (-8%)) compared with Day PT (*p*<0.0001, *d*=−1.903 and *p*<0.0001, *d*=0.574, respectively). METs on both Day PT-1 and Day PT+1 were also lower at T6 than at T0 (*p*<0.0001, *d*=1.016 and *p*<0.0001, *d*=0.574, respectively). Furthermore, at T6, METs were lower on Day PT-1 than on Day PT+1 (*p*<0.0001, *d*=−1.252), whereas no differences between days were observed at T0. Unlike the other PA measures, the 3-day average of METs was significantly lower at T6 than at T0 (-0.20 (-13%); *p*=0.003, *d*=0.536; Table S1).

Sedentary time (Table 2 and Figure 2F): Sedentary time was significantly longer on Day PT+1 than on Day PT at T0 (*p*=0.018; *d*=−0.509). No other differences were observed across days or time points. Similarly, the 3-day average of sedentary time did not differ between T0 and T6 (*p*=0.209; *d*=−0.216; Table S1).

#### Changes in clinical parameters

Following the 12-week intervention, significant changes were observed across several clinical parameters (see Table 3).

**Table 3.** Evolution of clinical parameters through the 12-week power-training intervention.

| Parameters | T0 | T12 | *p*-value |
| --- | --- | --- | --- |
| ***Strength and power*** |  |  |  |
| Upper-limb muscle strength (N) | **404.7 ± 79.4** | **418.5 ± 77.4** | **0.038** |
| Upper-limb muscle strength/BM (N/kg) | 5.22 ± 0.93 | 5.35 ± 0.85 | 0.307 |
| Maximal lower limb strength (N) | *134 ± 41* | *153 ± 52* | *0.099* |
| Muscle power (Takai; W) | **163 ± 32** | **175 ± 35** | **0.004** |
| ***Body composition parameters*** |  |  |  |
| Body mass (kg) | *78.1 ± 11.1* | *78.6 ± 10.4* | *0.063* |
| **- *Fat content by DXA*** |  |  |  |
| Upper limb fat mass (%) | *18.7 ± 5.9* | *18.2 ± 5.5* | *0.091* |
| Lower limb fat mass (%) | 21.6 ± 6.5 | 21.4 ± 6.5 | 0.271 |
| Trunk fat mass (%) | 32.7 ± 9.8 | 32.0 ± 9.3 | 0.109 |
| Android fat mass (%) | 38.0 ± 10.4 | 37.4 ± 9.4 | 0.127 |
| **Gynoid fat mass (%)** | *26.5 ± 6.8* | *25.8 ± 6.5* | *0.066* |
| Total fat mass (%) | *26.6 ± 7.4* | *26.0 ± 7.1* | *0.081* |
| **- *Lean content by DXA*** |  |  |  |
| Upper limb lean mass (kg) | **5.92 ± 0.91** | **6.10 ± 0.97** | **0.009** |
| Lower limb lean mass (kg) | **19.3 ± 2.42** | **19.8 ± 2.46** | **0.0001** |
| Appendicular lean mass (kg) | **25.2 ± 3.20** | **25.9 ± 3.31** | **<0.0001** |
| Gynoid lean mass (kg) | **8.08 ± 1.04** | **8.26 ± 0.99** | **0.002** |
| Total lean mass (kg) | **53.9 ± 5.77** | **54.8 ± 5.46** | **<0.0001** |
| **- *Thigh pQCT parameters*** |  |  |  |
| Muscle area (cm<sup>2</sup>) | **108 ± 17** | **113 ± 15** | **0.051** |
| Lean muscle area (cm<sup>2</sup>) | **102 ± 15** | **107 ± 14** | **0.036** |
| Intramuscular fat area (cm<sup>2</sup>) | 5.52 ± 2.96 | 5.52 ± 2.75 | 0.994 |
| Fat area (cm<sup>2</sup>) | 42.9 ± 16.3 | 43.5 ± 16.6 | 0.518 |
| Subcutaneous fat area (cm<sup>2</sup>) | 39.0 ± 16.1 | 39.3 ± 16.5 | 0.787 |
| **- *Vastus Lateralis - Ultrasound parameters*** |  |  |  |
| Muscle fascicle length (mm) | 10.8 ± 3.2 | 10.5 ± 2.6 | 0.707 |
| Muscle thickness (mm) | **1.67 ± 0.47** | **1.97 ± 0.38** | **0.023** |
| Muscle Pennation angle (°) | *12.8 ± 2.7* | *14.3 ± 2.7* | *0.079* |
| ***Physical performance parameters*** |  |  |  |
| 6-min walking distance (m) | **607 ± 94** | **634 ± 91** | **0.001** |
| Normal walking speed (4-m; m/s) | 1.47 ± 0.24 | 1.52 ± 0.21 | 0.132 |
| Usual TUG (3-m; s) | **9.71 ± 1.48** | **8.84 ± 1.30** | **0.001** |
| Fast TUG (3-m; s) | **6.62 ± 1.17** | **6.17 ± 1.05** | **0.0001** |
| Chair test (10 STS; s) | **20.4 ± 3.4** | **19.1 ± 3.6** | **0.026** |
| Stairs test (20s; n) | **32.3 ± 5.8** | **35.8 ± 6.2** | **<0.0001** |

Legend: Results are presented as mean ± SD. Paired-samples *t*-tests were performed, with $p < 0.05$ considered statistically significant (Bold) and $0.05 \leq p < 0.10$ considered indicative of a statistical tendency (Italics). Outcome abbreviations: TUG = Timed Up and Go; STS = sit-to-stand; BW = body weight; pQCT = peripheral quantitative computed tomography. Units: kg = kilograms; cm = centimeter; cm<sup>2</sup> = square centimeter; N = Newtons; % = percentage; m = meter; s = seconds; ° = degree; mm = millimeter.

### Strength and power

Upper-limb muscle strength increased significantly from 404.7±79.4 N at T0 to 418.5± 77.4 N at T12 (*p*=0.038; *d*=0.357), corresponding to a 3.3% increase. Estimated muscle power also increased significantly, from 163±32 W to 175±35 W (*p*=0.004; *d*=0.621), representing a 7.4% increase. In contrast, maximal lower limb strength increased from 134±41 N to 153±52 N, but this change did not reach statistical significance although there is a tendency toward significance (*p*=0.099; +14.2%). Similarly, relative upper-limb muscle strength did not significantly change (*p*=0.307).

### Body composition

Body composition analysis revealed significant increases in upper limbs [5.92±0.91 to 6.10±0.97 kg; *p*=0.009; *d*=0.455 (+3.0%)], lower limbs [19.3±2.4 to 19.8±2.4 kg; *p*=0.0001; *d*=0.702 (+2.6%)] and total [53.9±5.7 to 54.8±5.4 kg; *p*<0.0001; *d*=0.786 (+1.7%)] lean mass. Appendicular lean mass also increased from 25.2±3.1 to 25.9±3.3 kg [*p*<0.0001 (+2.8%)]. No significant changes were observed in body weight (78.1±11.1 to 78.6±10.4 kg; *p*=0.06 (+0.6%)) or fat mass parameters.

Regarding muscle composition, lean muscle area increased significantly from 102±15 to 107±14 cm² [*p*=0.036; *d*=0.434 (+4.9%)] whereas total muscle area tend to increase from 108±17 to 113±15 cm² [*p*=0.051 (+4.6%)]. For muscle architecture, muscle thickness increased significantly from 1.67±0.47 to 1.97±0.38 mm [*p*=0.023; *d*=0.497 (+18.0%)], while muscle pennation angle showed a tendency toward an increase [12.8±2.7 to 14.3±2.7 °; *p*=0.08 (+11.7%)].

### Physical performance

Six-minute walking distance increased from 607±94 to 634±91 m [*p*=0.001; *d*=0.639 (+4.4%)]. Usual TUG time [from 9.71±1.48 to 8.84±0.30 s; *p*=0.001; (improvement:9.0%)] and fast TUG time [from 6.62±1.17 to 6.17±1.05 s; *p*=0.0001; *d*=−0.719 (improvement: 6.8%)] decrease significantly. Chair test performance [from 20.4±3.4 to 19.1±3.6 s; *p*=0.026; *d*=−0.498 (improvement:6.4%)] and stair test performance [from 32.3±5.8 to 35.8±6.2 repetitions; *p*<0.0001; *d*=1.044 (improvement:10.8%)] improved. Normal walking speed didn’t change statistically (from 1.47±0.24 to 1.52± 0.21 m/s, *p*=0.132).

#### Associations between physical activity behavior parameters and clinical parameters

Some correlations between changes in physical activity behavior parameters and changes in clinical parameters were observed following the 12-week PT intervention. Day PT-1 was selected for the association analyses because it was the only non-training day showing significant changes in PA behavior between T0 and T6, whereas no significant T0-T6 changes were observed on Day PT+1. Changes in PA time between Day PT-1 and Day PT at T6 were positively correlated with changes in Fast TUG time (r=0.40, *p*=0.02; Figure 3A) and muscle pennation angle (r=0.47, *p*=0.02; Figure 3D). The increase in Fast TUG time signifies slower speed and thus, the positive correlation indicates that an increase in PA time leads to a poorer performance in Fast TUG. Changes in estimated muscle power were negatively correlated with changes in number of steps between Day PT-1 and Day PT at T6 (r=−0.39, *p*=0.02; Figure 3B), indicating that greater increases in daily steps were associated with smaller increases in estimated muscle power. Changes in upper limb strength were also positively correlated with changes in number of steps (r=0.34, *p*=0.04; Figure 3C). All correlations results are presented in Table S2.

**Figure 3.**
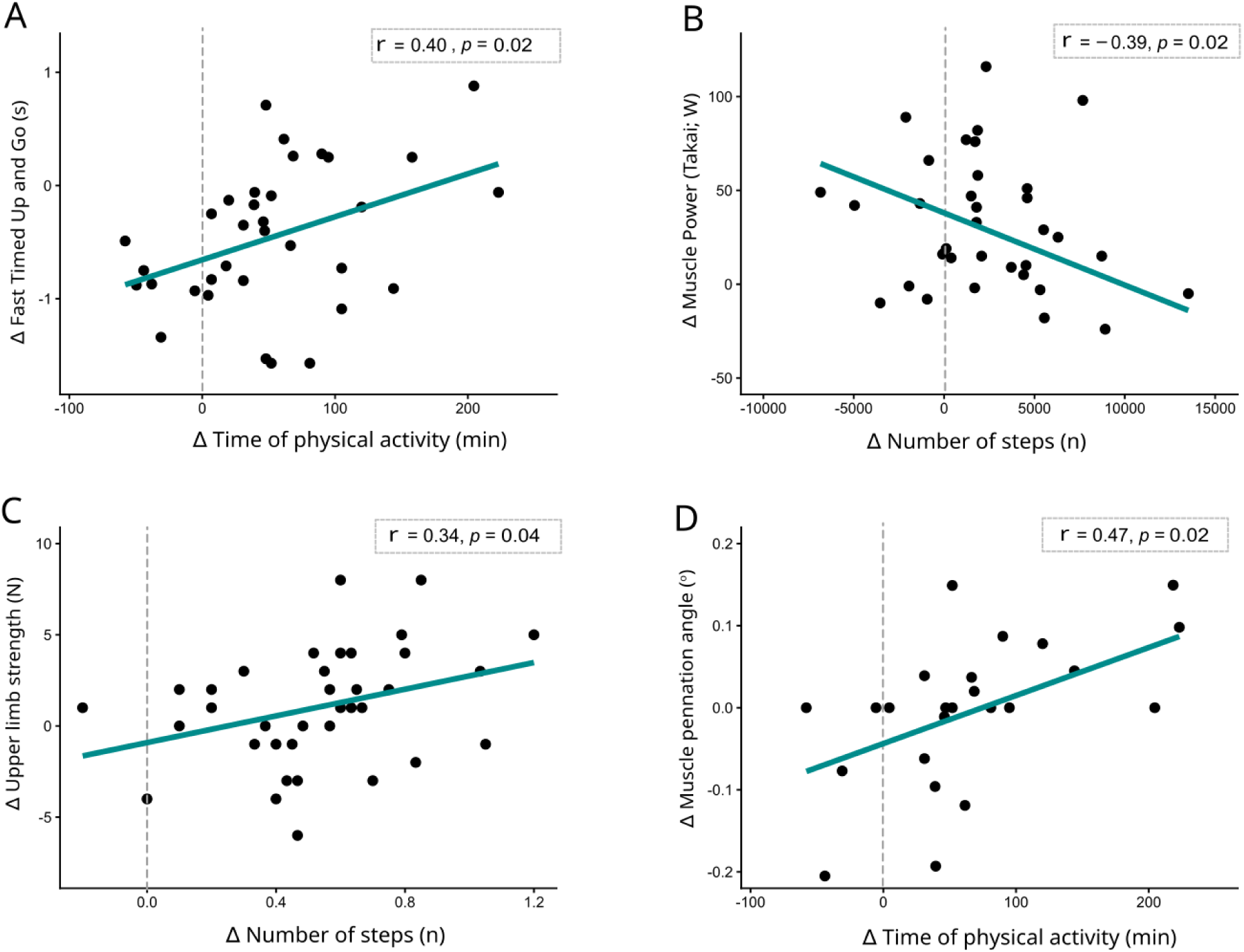
Correlations between physical activity parameters changing significantly at mid-intervention and clinical parameters changing significantly following the 12-week PT intervention. Legend: Δ changes of the physical activity parameters were calculated at mid-intervention (Day PT - Day PT-1). Δ changes of the clinical parameters were calculated after the 12-week intervention (Post-Pre). (A) Δ fast - Timed Up and Go (s) and Δ Time of physical activity (min); (B) Δ Takai [(W; height (cm)/10-s STS repetitions)]) and Δ number of steps (n); (C) Δ Upper limb strength (kg) and Δ Number of steps (n); (D) Δ Muscle pennation angle (°) and Δ Time of physical activity (min). r = Pearson’s correlation coefficient. The black dots represent the individual data, and the blue line refers to the linear regression line.

## 4. Discussion

The present study aimed to examine the day-to-day variability of physical activity (PA) behavior surrounding training sessions, and to explore its possible associations with clinical adaptations following a 12-week PT intervention in older men. First, the findings highlight the importance of considering day-to-day variability in PA behavior, as PA was significantly modulated on the days preceding and following training sessions between pre- and mid-intervention, whereas no change was detected when PA was averaged across days. Second, changes in some PA behavior at mid-intervention were associated with some clinical adaptations. Together, these results suggest that PA behavior surrounding training sessions should be considered as an important compensatory factor contributing to variability in training adaptations. Specifically, adaptation may depend not only on the training stimulus but also on how older men adjust their spontaneous activity before and after each session.

The behavioral pattern change observed at mid-intervention could be consistent with activity compensation, where individuals subconsciously adjust their spontaneous physical activity in response to structured exercise (Gray et al., 2018). In the present context, the reduction in activity on Day PT-1 may reflect anticipatory regulation, whereas the decrease on Day PT+1 may indicate residual fatigue or recovery-related processes following the training session (Wender et al., 2022). Given that power training involves high-velocity contractions and neuromuscular demands (Porter & Metabolism, 2006), such sessions may induce transient fatigue, leading to reduced voluntary activity in the subsequent 24 hours (Egerton et al., 2016). Rather than adding the training stimulus to usual activity, participants may therefore have redistributed their daily energy expenditure. The greater reduction on Day PT-1 may reflect self-regulation to conserve energy and optimize performance, whereas the smaller reduction on Day PT+1 may reflect recovery from the preceding session, potentially due to muscle soreness or fatigue (Harada, 2022). This pattern may be particularly relevant in older adults, who typically have slower recovery process and may require additional time to recover from exercise-induced physiological stress. Given that training sessions were performed every two days, the intervening day may have provided an important opportunity for recovery and contributed to the observed redistribution of daily activity.

Moreover, no significant differences were observed between baseline and mid-intervention when PA variables were averaged across the three monitored days except for METs. This finding highlights that aggregated measures may obscure meaningful behavioral adaptations (Bussmann & van den Berg -Emons, 2013). Although multi-day or weekly averages are commonly used to characterize PA behavior (Migueles et al., 2017) they may overlook transient but systematic changes occurring in relation to exercise sessions. Consequently, null findings based on aggregated measures may not necessarily indicate an absence of behavioral change. Assessing PA at a finer temporal resolution and in relation to exercise timing may provide a more accurate representation of behavioral adaptations and improve comparability across intervention studies though standardized approaches (Welk et al., 2019). Therefore, future studies should consider not only the total PA volume but also its temporal organization and relationship with exercise exposure.

Furthermore, a novel aspect of this study lies in the associations observed between changes in PA behavior and clinical adaptations. Although substantial inter-individual variability in response to exercise training is well documented (Bouchard et al., 2001), its underlying determinants remain incompletely understood. Traditionally, variability in adaptation has been attributed to factors such as genetics, baseline fitness, age, and differences in training responsiveness (Mann et al., 2014; Noone et al., 2024). The present findings suggest that PA behaviors occurring outside structured training exercise may also contribute to this heterogeneity. Each training stimulus occurs within a broader behavioral context characterized by fluctuations in PA and sedentary behavior, suggesting that exercise adaptation may reflect the interaction between structured training and daily PA. Continuous PA monitoring could therefore provide additional insight into individual differences in training responses and support the development of more individualized exercise strategies.

Greater fluctuations in activity should not necessarily be interpreted as evidence of greater adaptation, as larger changes around training sessions may instead reflect physiological perturbation, recovery demands, or fatigue (Enoka & Duchateau, 2016; Riou et al., 2019). Conversely, reducing spontaneous activity may decrease the additional mechanical and metabolic stress accumulated outside training, potentially supporting recovery and adaptation (Kellmann et al., 2018). The divergent associations across outcomes further suggest that changes in daily activity may influence different adaptations through distinct mechanisms. For example, habitual movement may provide additional mechanical stimuli relevant to muscle architecture, consistent with evidence linking habitual loading to skeletal muscle and bone characteristics in older adults (Wullems et al., 2024).

Several associations observed in Table S2 further illustrate the complex relationship between daily PA behavior and training adaptations. Greater PA time was associated with lower fast TUG speed, which may indicate that individuals who accumulated more daily activity also exhibited slower fast-paced mobility performance following the intervention. Although counterintuitive, this association may reflect differences in how daily activity was accumulated rather than a simple beneficial or detrimental effect of greater activity. Similarly, a higher number of daily number of steps was associated with lower muscle power. One possible explanation is that step count captures the quantity of ambulatory activity but does not characterize its intensity, mechanical demands, or movement characteristics. Therefore, a greater number of steps may not necessarily provide the type or magnitude of mechanical stimulus required to improve muscle power. In contrast, the positive association between step count and upper limb strength should be interpreted cautiously, as it may represent a coincidental association rather than a biologically meaningful relationship. Also, some additional associations reached a statistical tendency and may nevertheless warrant consideration. For example, the tendency for greater physical activity time to be associated with greater muscle power may be consistent with the possibility that habitual activity contributes to the maintenance or development of neuromuscular capacity. However, given the exploratory nature of these analyses and the number of associations examined, such findings should be interpreted cautiously and considered hypothesis-generating rather than confirmatory. Future studies should therefore incorporate direct measures of fatigue, muscle damage, recovery, and neuromuscular function to determine how structured exercise, daily activity behavior, and recovery interact to influence training responses.

Some limitations should be acknowledged. First, the study included only older men, which limits the generalizability of the findings to women, less functional older adults, and other populations. Second, physical activity was not assessed at T12 and hence it remains unclear whether the behavioral changes observed at T6 were maintained, attenuated, or further modified during the remainder of the intervention. Third, although accelerometry provided objective measures of physical activity, it did not capture contextual information or distinguish between different types of activities. Fourth, the observational nature of the correlation analyses precludes causal inferences regarding the relationship between activity behavior and physiological adaptations. Moreover, given the a posteriori per-protocol design, these findings should be considered exploratory. Replication in larger and more diverse cohorts, including less functional individuals, with assessments throughout the intervention is warranted. Future studies should also investigate physiological markers of fatigue, recovery, and neuromuscular function underlying the relationship between behavioral regulation and exercise adaptations.

## 5. Conclusion

The current study demonstrates that daily physical activity behavior is modulated during a 12- week power training intervention in older men, with distinct fluctuations observed around training sessions at mid-intervention. Furthermore, the observed associations between changes in PA behavior and clinical adaptations suggest that habitual movement patterns may represent an additional behavioral factor contributing to inter-individual variability in training responses. Although the underlying mechanisms remain to be elucidated, these findings emphasize that exercise-induced adaptations may not solely depend on the prescribed training stimulus but may also be influenced by how individuals regulate their daily physical activity outside structured exercise sessions. Future research incorporating continuous behavioral monitoring alongside markers may help clarify the role of daily activity regulation in optimizing individualized exercise interventions.

## Acknowledgement

We sincerely thank all the participants and all the trainees for their commitment and contribution to this study.

## CRediT authorship contribution statement

**Layale Youssef:** Formal analysis, Data curation, Writing-Original draft. **Haroun El-Oueslati**: Writing-Review & Editing. **Justine Persouyré:** Data curation, Writing-Review & Editing. **Charlotte Pion:** Investigation, Writing-Review & Editing. **Paula Lago:** Writing-Review & Editing. **Marc Bélanger:** Conceptualization, Methodology, Visualization, Writing-Review & Editing. **Mylène Aubertin-Leheudre**: Conceptualization, Methodology, Visualization, Supervision, Writing-Review & Editing.

---

**Table S1.** Average daily physical activity parameters across the three days surrounding the power training session at baseline (T0) and mid-intervention (T6).

| Parameters | All 3 days at T0 | All 3 days at T6 | *p-values*<br>*(3-days at T0 vs. 3-days at T6)* |
| --- | --- | --- | --- |
| Total PA (min/d) | 118 ± 98 | 106 ± 54 | 0.513 |
| Number of steps (n/d) | 8729 ± 3026 | 8381 ± 2567 | 0.451 |
| TEE (kcal/d) | 2645 ± 753 | 2436 ± 413 | 0.103 |
| AEE(kcal/d) | 684 ± 829 | 558 ± 365 | 0.434 |
| METs (Mets/d) | **1.44 ± 0.34** | **1.25 ± 0.18** | **0.003** |
| SED time (h/d) | 7.77 ± 2.09 | 8.04 ± 2.16 | 0.209 |

Legend: Results are presented as mean ± SD. T0 = baseline (pre-intervention); T6 = mid-intervention (6 weeks). All 3 days refer to the power training day and the days before and after the power training day. min = minutes; kcal = kilocalories; METs = metabolic equivalent tasks; h = hours; PA: physical activity; TEE: total energy expenditure; AEE: active energy expenditure. Physical behavior parameters were assessed using a validated 3D accelerometer (armband Sensewear®). Statistical analyses were performed using paired-samples t-tests with a *p*-value < 0.05 considered statistically significant.

**Table S2.** Associations between physical activity parameters changing significantly around the power training session at mid-intervention and clinical parameters changing significantly following the 12-week intervention.

| Clinical parameter | PA time |  | Steps number |  | TEE |  | AEE |  | METs |  | Sedentary time |  |
| --- | --- | --- | --- | --- | --- | --- | --- | --- | --- | --- | --- | --- |
|  | r | *p* | r | *p* | r | *p* | r | *p* | r | *p* | r | *p* |
| 3-m normal TUG (s) | 0.22 | 0.21 | 0.22 | 0.20 | 0.04 | 0.81 | 0.18 | 0.30 | 0.07 | 0.69 | 0.00 | 0.99 |
| 3-m fast TUG (s) | **0.40** | **0.02** | 0.13 | 0.46 | 0.06 | 0.73 | **0.34** | **0.04** | 0.20 | 0.24 | -0.07 | 0.67 |
| Chair test (10 Sit-to-Stand; s) | 0.03 | 0.89 | 0.00 | 0.98 | 0.28 | 0.19 | 0.09 | 0.68 | 0.01 | 0.95 | 0.01 | 0.96 |
| Stairs test (20s; n) | 0.01 | 0.93 | -0.03 | 0.86 | 0.01 | 0.94 | -0.01 | 0.97 | -0.18 | 0.31 | -0.2 | 0.24 |
| Upper limb strength (N) | -0.21 | 0.23 | **0.34** | **0.04** | -0.22 | 0.20 | -0.19 | 0.26 | -0.10 | 0.55 | -0.15 | 0.38 |
| 6 min walking distance (m) | 0.09 | 0.61 | 0.28 | 0.09 | 0.03 | 0.85 | 0.04 | 0.81 | 0.15 | 0.38 | 0.14 | 0.42 |
| Muscle power (Takai; W) | -0.33 | 0.05 | **-0.39** | **0.02** | -0.10 | 0.58 | -0.32 | 0.06 | -0.12 | 0.49 | 0.33 | 0.06 |
| Muscle thickness (mm) | -0.07 | 0.75 | 0.00 | 0.99 | -0.38 | 0.06 | -0.14 | 0.52 | 0.03 | 0.88 | -0.2 | 0.35 |
| Muscle pennation angle (°) | **0.47** | **0.02** | 0.36 | 0.08 | 0.23 | 0.28 | **0.41** | **0.04** | 0.37 | 0.07 | -0.3 | 0.16 |
| Upper limb lean mass (kg) | 0.08 | 0.64 | -0.06 | 0.74 | 0.02 | 0.92 | 0.11 | 0.51 | 0.10 | 0.56 | -0.02 | 0.89 |
| Lower limb lean mass (kg) | 0.04 | 0.82 | 0.14 | 0.42 | 0.08 | 0.67 | 0.04 | 0.82 | 0.16 | 0.36 | -0.02 | 0.91 |
| Gynoid lean mass (kg) | 0.05 | 0.76 | -0.06 | 0.74 | 0.1 | 0.57 | 0.01 | 0.96 | -0.14 | 0.41 | 0.33 | 0.05 |
| Total lean mass (kg) | -0.05 | 0.79 | -0.10 | 0.57 | -0.04 | 0.80 | -0.21 | 0.21 | -0.05 | 0.76 | 0.3 | 0.07 |
| Appendicular lean mass (kg) | -0.17 | 0.34 | -0.16 | 0.34 | -0.18 | 0.29 | -0.24 | 0.16 | -0.11 | 0.52 | 0.14 | 0.42 |
| Muscle area (cm<sup>2</sup>) | 0.32 | 0.12 | 0.29 | 0.17 | 0.24 | 0.26 | 0.40 | 0.09 | 0.34 | 0.11 | -0.35 | 0.09 |
| Lean muscle area (cm<sup>2</sup>) | -0.21 | 0.31 | -0.38 | 0.06 | -0.36 | 0.06 | -0.25 | 0.21 | -0.14 | 0.48 | -0.14 | 0.50 |

Legend: Values are Pearson correlation coefficients (r) and corresponding *p*-values. PA = physical activity; TEE = total energy expenditure; AEE = active energy expenditure; METs = metabolic equivalent tasks. Significant associations ( $p < 0.05$ ) are shown in bold, and tendencies toward an association ( $0.05 \leq p < 0.10$ ) are shown in italics.

